# *Bmmp* influences wing shape by regulating anterior-posterior and proximal-distal axis development

**DOI:** 10.1101/2021.07.26.453796

**Authors:** Yunlong Zou, Xin Ding, Li Zhang, Lifeng Xu, Shubo Liang, Hai Hu, Fangyin Dai, Xiaoling Tong

## Abstract

Insect wings are subject to strong selective pressure, resulting in the evolution of remarkably diverse wing shapes that largely determine flight capacity. However, the genetic basis and regulatory mechanisms underlying wing shape development are not well understood. The silkworm *Bombyx mori micropterous* (*mp*) mutant exhibits shortened wing length and enlarged vein spacings, albeit without changes in total wing area. Thus, the *mp* mutant comprises a valuable genetic resource for studying wing shape development. In this study, we used molecular mapping to identify the gene responsible for the *mp* phenotype and designated it *Bmmp*. Phenotype-causing mutations were identified as indels and single nucleotide polymorphisms in non-coding regions. These mutations resulted in decreased *Bmmp* mRNA levels and changes in transcript isoform composition. *Bmmp* null mutants were generated by CRISPR/Cas9 and exhibited significantly smaller wings. By examining the expression of genes critical to wing development in wildtype and *Bmmp* null mutants, we found that *Bmmp* exerts its function by coordinately modulating anterior-posterior and proximal-distal axis development. We also studied a *Drosophila mp* mutant and found that *Bmmp* is functionally conserved in *Drosophila*. The *Drosophila mp* mutant strain exhibits curly wings of reduced size and a complete loss of flight capacity. Our results increase our understanding of the mechanisms underpinning insect wing development and reveal potential targets for pest control.

## Introduction

Wings endow insects with tremendous adaptive advantages because they enhance survival and fitness by making it possible to change environments rapidly. Insect wings are constantly subject to adaptive evolution and exhibit remarkable diversity in shape. Changes in wing shape result in differences in flight capacity, leading to variations in insect lifestyle [1, 2]. For example, dimorphism in wing shape occurs in a wide range in insects, such as rice planthoppers [3, 4] and aphids [5]. Long-winged morphs can fly, which allows them to escape adverse habitats and track changing resources, whereas short-winged morphs are flightless, but usually possess higher fecundity [1, 2]. In the order Lepidoptera, wing shapes are distinctly different between migratory species and non-migratory species. Typically, migratory moths and butterflies have relatively narrower forewings with straighter costal margins compared to those of non-migratory species [6].

Coordinated regulation of anterior-posterior (A-P) and proximal-distal (P-D) wing axis development plays a crucial role in correct wing shape formation. During this process, wing patterning and proliferation are coordinately modulated by relay signals [7]. For A-P axis development, posterior compartment identity is specified by the *engrailed* gene, which then activates the expression of *hedgehog* [7]. The secreted Hedgehog protein traverses the A-P border and induces expression of *Dpp* and *Wnt1* in anterior cells close to the border [8]. During the process of wing P-D axis development, *apterous* is expressed in the dorsal compartment and activates Notch signaling, which in turn induces *Wnt1* activity at the dorsal-ventral (D-V) border [9, 10]. Wnt1 helps establish the P-D axis of the wing by activating the *Distal*-*less* gene, which specifies the most distal regions of the wing [11, 12]. Reduced levels of Dpp affect both the width and length of the resulting wing and significantly decrease total wing area [13]. Moderate and uniform amounts of exogenous Wnt1 stimulate proliferative wing growth, leading to enlargement of the prospective wing [14].

The identification of new factors that influence wing shape will expand our understanding of the genetic basis of wing diversity. We hypothesized that as-yet uncharacterized key regulators coordinately regulate both A-P and P-D axis signals during wing development. To examine this developmental process more closely, we used the silkworm *Bombyx mori* (Lepidoptera, Bombycidae) *micropterous* (*mp*) mutant, which exhibits shortened wing length and enlarged vein spacings. We identified the gene responsible for the *mp* phenotype and designated it *Bmmp*. We found mutations in the noncoding regions of *Bmmp* that result in decreased *Bmmp* mRNA levels and changes in transcript isoform composition. In addition, we generated a *Bmmp* null mutant and determined that *Bmmp* exerts its effect on wing shape by regulating wing A-P and P-D axis development.

## Results

### Characterization of the silkworm *micropterous* (*mp*) mutant wing phenotype

To characterize the wing phenotype of the silkworm *mp* mutant, we compared pupae and moth wing phenotypes of *mp* and *Dazao* wildtype (WT) silkworms. Whereas the wings of WT pupae fully cover the third abdominal segment, the wings of *mp* pupae only cover the second abdominal segment, leaving the third abdominal segment naked (**Fig 1A**). Further examination demonstrated that the wing length of *mp* moths was significantly shorter than that of WT moths within each sex, although there was not a significant difference in total wing area (**Fig 1B, 1C, 1D**). In addition, there was significantly greater spacing between adjacent longitudinal veins in the wings of *mp* moths compared to those of WT moths within each sex (**Fig 1E**). These results demonstrate that the *mp* phenotype is not associated with a specific gender. To reflect the overall changes in wing morphology, we divided the wing length by the sum of longitudinal veins spacings. The resulting value is significantly smaller for *mp* moths than for WT moths within each sex (**Fig 1F**).

**Figure 1.**
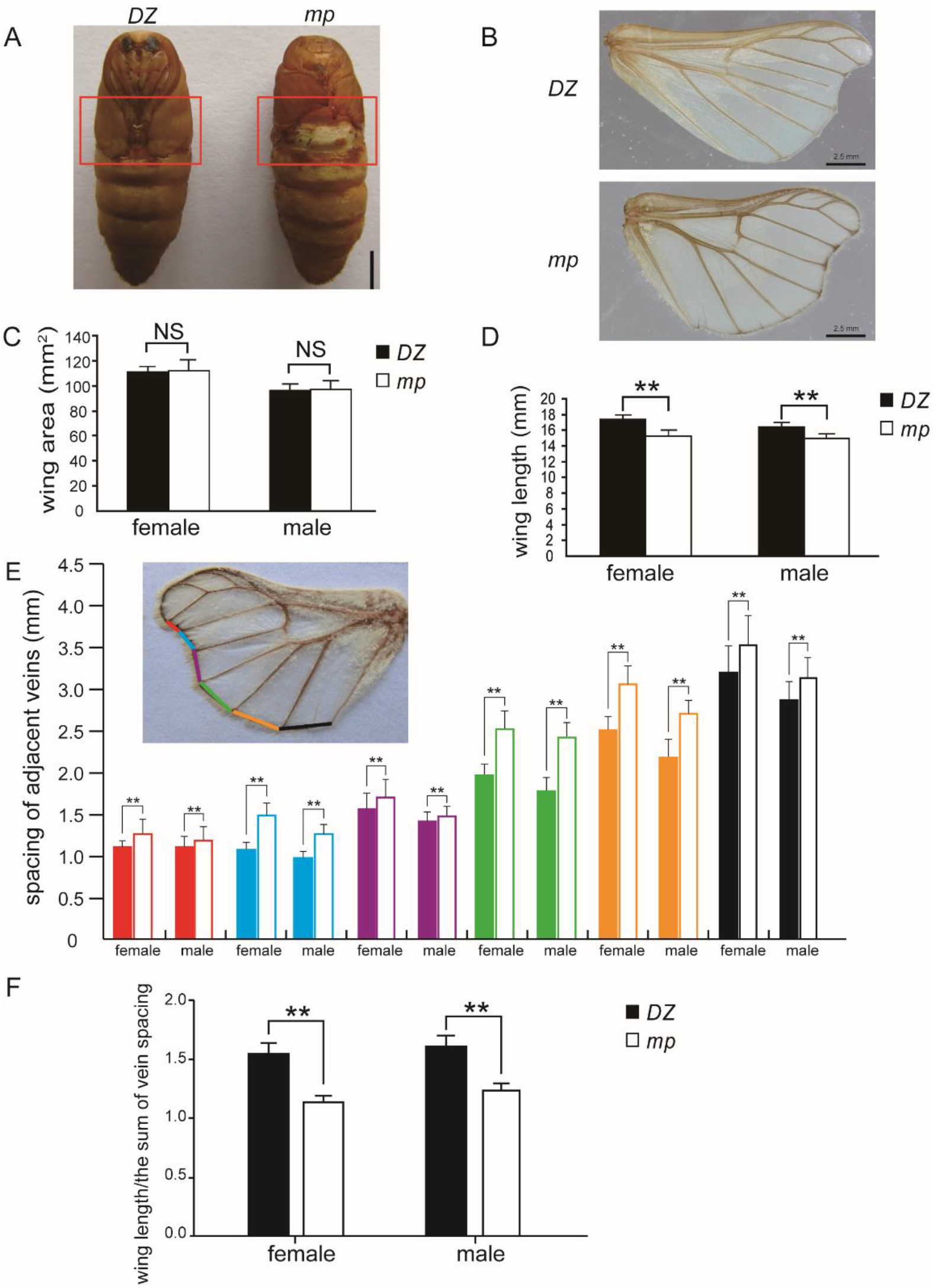
Characterization of wing phenotypes of silkworm *mp* mutant. (A) Representative photograph of male WT (left) and *mp* (right) silkworm pupae. Compared to WT pupae, *mp* pupae exhibit a naked third abdominal segment in the pupal stage, suggestive of a modified wing phenotype. The phenotypically variable region is highlighted within the red rectangular frame. Female *mp* silkworms exhibit the same phenotype. Scale bar, 5 mm. (B) Representative photograph of female WT (top) and *mp* (bottom) silkworm moth wings showing the differences in shape. Male *mp* silkworms exhibited identical phenotypes. Note that scale hairs were removed from the wing to exhibit wing shape characteristics more clearly. Scale bar, 2.5 mm. (C-F) Wings of both male and female WT and *mp* silkworm moths were measured using ImageJ following imaging with a digital microscope. (C) Within each sex, total wing area of WT and *mp* moths do not differ significantly. (D) Wing length was significantly shorter in *mp* moths compared to WT moths. (E) The spacings between adjacent longitudinal veins were significantly larger in *mp* moths compared to WT moths. Filled columns, WT; unfilled columns, *mp*. Column colors correspond to specific vein spacings indicated by lines of the same color overlayed on the photographed wing. (F) Wing length was divided by the sum of spacings between longitudinal veins to reflect overall wing shape. The resulting values were significantly different between *mp* and WT moths. N = 30 for both male and female *mp* and *Dazao* silkworm moths. **, P < 0.01; NS, not significant. *DZ*, *Dazao*, used as wildtype control; *mp*, silkworm *mp* mutant. Error bars represent SD.

### Molecular mapping and analysis of candidate genes responsible for the *mp* phenotype

To identify candidate gene(s) responsible for the *mp* phenotype, we performed a genetic linkage analysis using *B. mori* simple sequence repeat (SSR) markers and newly designed markers polymorphic between WT and *mp* silkworms. Initially, we roughly mapped the *mp* phenotype using 456 BC_1_M individuals and SSR markers on the eleventh linkage group. The results indicate that the gene responsible for the *mp* phenotype is located within a 12.1-cM region linked to SSR marker S1146 (**Fig 2A and 2B**). Subsequent fine mapping with 320 BC_1_M and newly designed primer sets narrowed the *mp* locus to an approximately 260-kb region between markers 2810A and 2810C on the nscaf2810 scaffold. The 2810M marker was tightly linked with the *mp* locus (**Fig 2C**). Two candidate genes (*KWMTBOMO06923* and *KWMTBOMO06924*) were identified within the 260 kb region, based on annotated sequences obtained from the SilkBase database [15] (**Fig 2D**).

**Figure 2.**
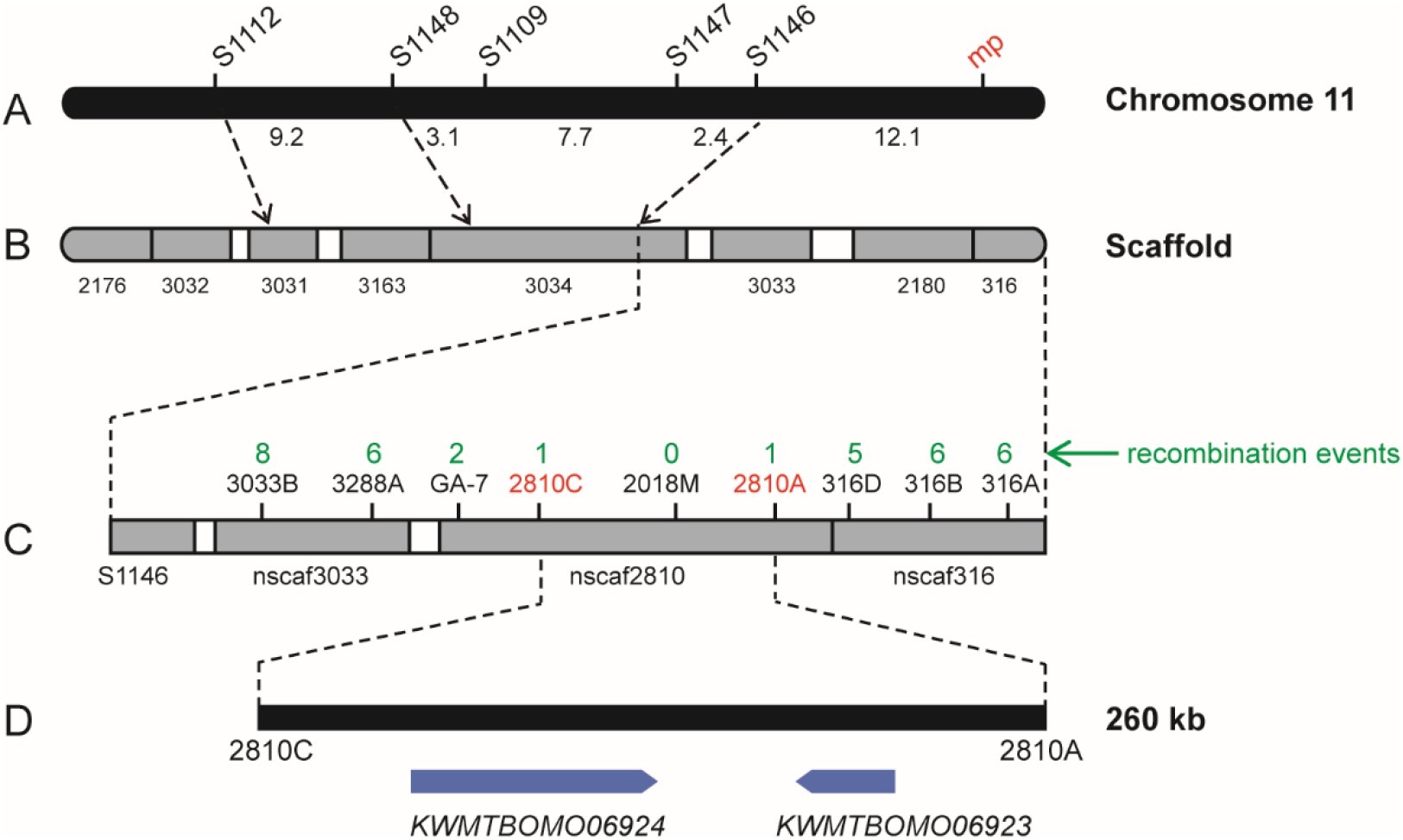
Molecular mapping of candidate genes responsible for the *mp* phenotype. (A) Map of *B. mori* chromosome 11 with locations of SSR markers used in this study. Five SSR markers and the *mp* locus are labeled above the map. Map distances are shown in cM. (B) Schematic of scaffolds on chromosome 11. Gray boxes represent the assembled scaffolds; their respective serial numbers are shown below. S1112 mapped to nscaf3031, whereas S1148, S1109, S1147, and S1146 all mapped to nscaf3034. (C) Expanded view of genomic scaffolds used for fine mapping of the *mp* locus. Newly designed primer sets are shown on the map, and the numbers above them indicate the respective recombination events in 320 BC_1_M progeny. The *mp* locus was tightly linked to 2810M, located between markers 2810A and 2810C. (D) Gene annotation in the *mp* linked region. Two genes were predicted in this region, namely *KWMTBOMO06923* and *KWMTBOMO06924*.

Because the functions of *KWMTBOMO06923* and *KWMTBOMO06924* are uncharacterized, we searched for mutations responsible for the *mp* phenotype by comparing the corresponding genomic sequences from *mp* silkworms and silkworms with normal wings. Although synonymous single nucleotide polymorphisms (SNPs) mutations were identified in the *mp KWMTBOMO06923* and *KWMTBOMO06924* genes, no mutations that changed the sequences of the predicted translated proteins were found. We next surveyed all introns within *KWMTBOMO06924*, as well as putative regulatory regions 2 kb upstream and downstream from the gene. A total of 59 indels and 101 SNPs specific to the *mp* mutant were identified. (**Table S1 and Table S2**).

Multiple transcript isoforms of *KWMTBOMO06924* are annotated in the *B. mori* EST database (http://sgp.dna.affrc.go.jp/KAIKObase/). To obtain more detailed isoform information, we generated and sequenced *KWMTBOMO06924* cDNA libraries. Sequence alignments revealed that the *KWMTBOMO06924* gene is comprised of 10 exons, spanning 36.79 kb of genomic DNA. A total of 28 *KWMTBOMO06924* transcript isoforms were identified in WT wing discs. The full-length cDNA sequence contained a 1443-bp open reading frame encoding 481 amino acids, consistent with the cDNA clone (fwd-02K11) retrieved from the *B. mori* EST database. The protein encoded by the full-length transcript isoform contains three functional domains (BTB, BACK and TLDc) as determined using the SMART online prediction tool. We next examined *KWMTBOMO06924* transcript isoform composition and expression in wing discs from *mp* and WT silkworms. Of the 28 transcript isoforms detected in the WT strain, only 6 were recovered in the *mp* mutant. In addition, one unique transcript was identified in the *mp* silkworms (**Fig 3**). An intact BTB domain, encoded by exons 1-3, was present in all transcript isoforms identified in both silkworm strains. Quantitative RT-PCR analysis of wing discs from silkworms at the initiation of the wandering stage revealed that total *KWMTBOMO06924* mRNA levels were significantly lower in *mp* vs. WT silkworms during this critical period of wing development (**Fig 4**).

**Figure 3.**
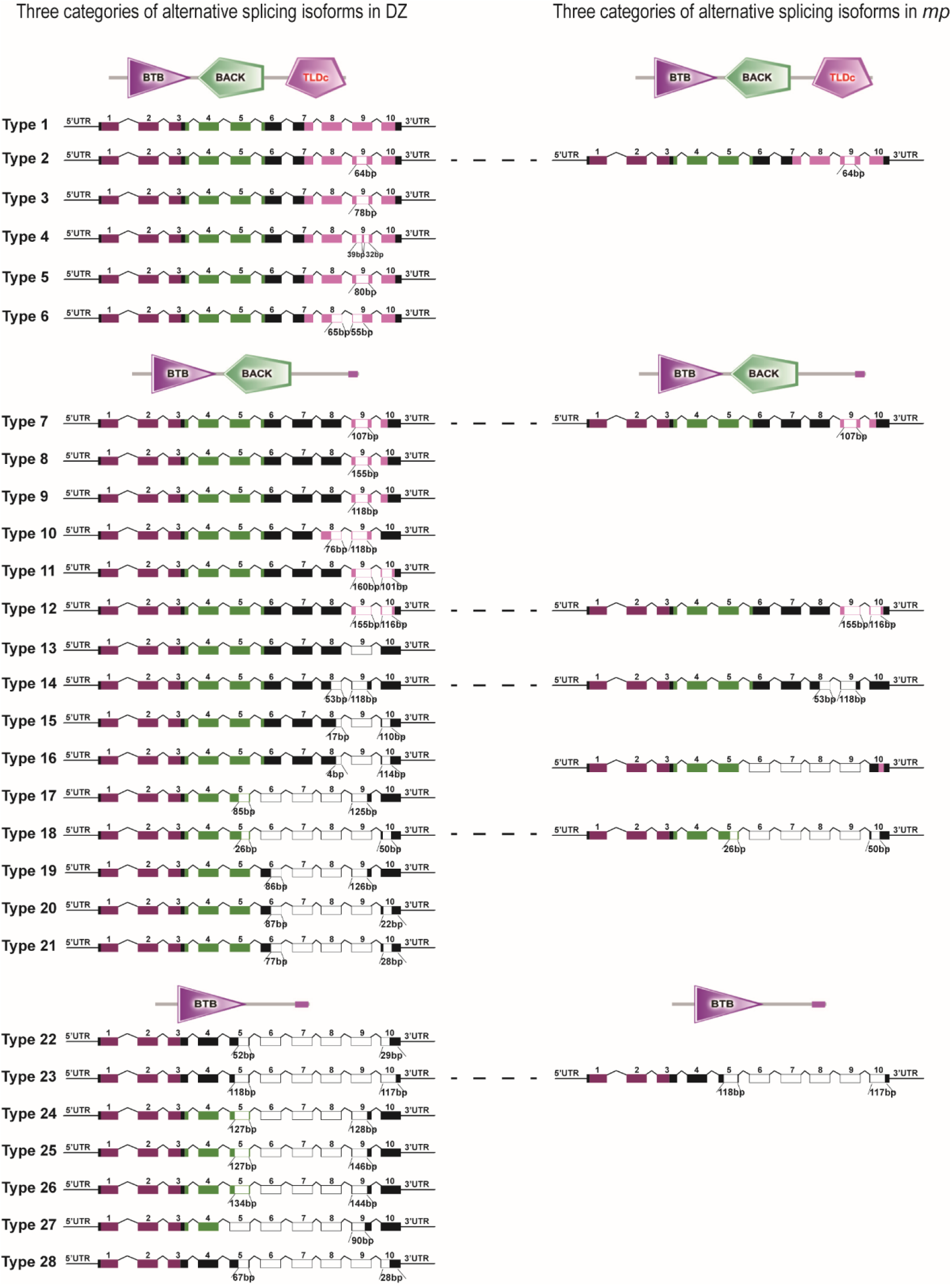
Transcript isoforms in wing discs from *mp* and WT silkworms. Twenty-eight distinct transcript isoforms were identified in wing discs from WT silkworms. Six of the 28 transcript isoforms identified in WT silkworms were also recovered in wing discs from *mp* silkworms, plus one unique isoform. Dotted lines indicate transcript isoforms identified in both WT and *mp* silkworms. Exon box colors correspond to the encoded domains indicated above each category. Unfilled regions represent exons and portions of exons not included in the specific mature mRNA transcript shown in the figure. Sizes (nucleotides) of truncated exons are shown below the truncations. *DZ*, *Dazao*, used as wildtype control; *mp*, silkworm *mp* mutant.

**Figure 4.**
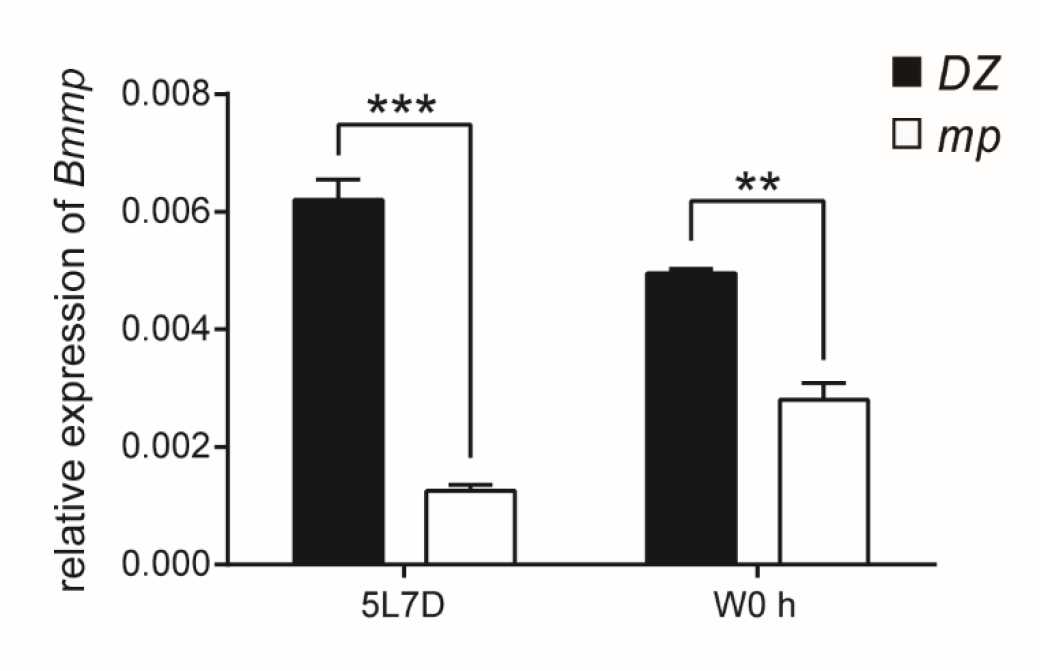
Relative *Bmmp* mRNA levels in wing discs from WT and *mp* silkworms. *Bmmp* mRNA levels in wing discs from WT and *mp* silkworms were quantified by qRT-PCR. Relative *Bmmp* mRNA levels were significantly higher in WT silkworms compared to *mp* silkworms at two different developmental stages (5L7D and W0 h). 5L7D, fifth instar at day 7; W0 h, wandering stage at 0 h; ***, P < 0.001; **, P < 0.01. *DZ*, *Dazao,* used as wildtype control; *mp*, silkworm *mp* mutant. N=3 for both *Dazao* and *mp* silkworms at 5L7D and W0 h, respectively. Values are relative to expression of eukaryotic translation initiation factor 4A (defined as 1). Experiment was independently repeated three times. Error bars represent SEM.

Together, these results suggest that *KWMTBOMO06924* is responsible for the *mp* phenotype, with causative mutations localized to regulatory regions. Thus, we designated this gene *Bmmp*. The results are consistent with two possible causes for the *mp* phenotype: (1) decreased total *Bmmp* mRNA levels, and (2) reduced variation of *Bmmp* transcript isoforms.

### Null mutation of *Bmmp* results in a significant reduction in wing size

To elucidate the function of the *Bmmp* gene in wing development and morphology, we utilized the CRISPR/Cas9 system to disrupt *Bmmp*. We selected four genomic targets spanning 130 bp in exon 1 to generate large fragment deletions (**Fig 5A**). Since this region is shared across isoforms, any frame-shift mutations would be predicted to abolish all functional transcripts. sgRNAs were synthesized *in vitro* for the genomic targets, mixed with Cas9 protein, and injected into the preblastoderm of *Dazao* embryos. In total, 110 injected embryos hatched, and 81 individuals survived to an adult stage. Out of these 81 silkworms, 67 exhibited markedly smaller wings in pupal and adult stages, compared to uninjected WT controls (**Fig 5B and 5C**). To confirm that the *Bmmp* deletions caused the decrease in wing size, genomic DNA was extracted from three moths with small wings. Regions spanning the four sgRNA targets were amplified by PCR, subcloned, and sequenced. As expected, the three selected moths contained *Bmmp* deletions and no wildtype sequences (**Fig S1**). Notably, five distinct mutations were identified in moth #11 (**Fig S1**), demonstrating the presence of mosaicism in silkworms of the injected generation (generation 0, G_0_).

**Figure 5.**
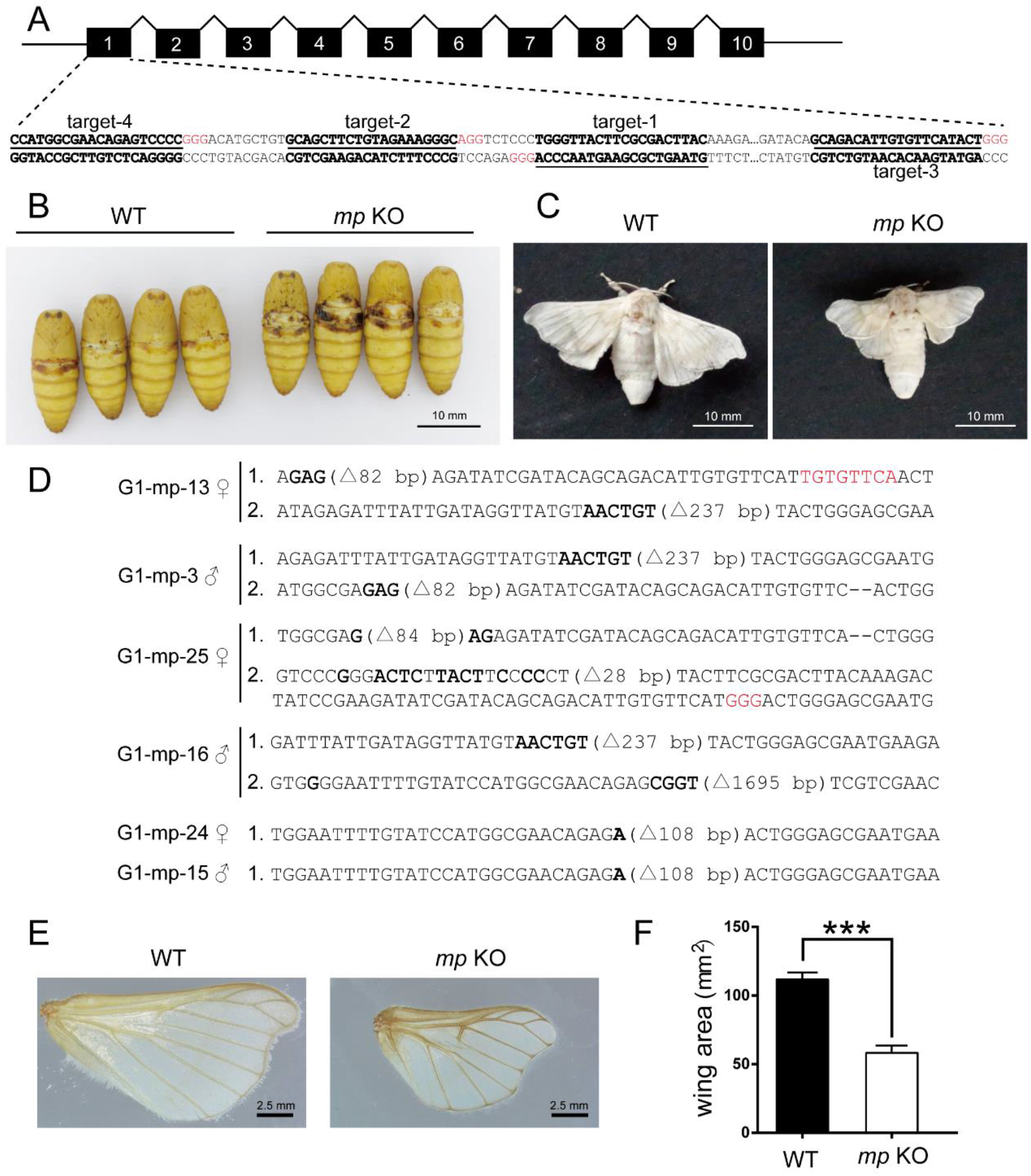
Construction of *Bmmp* knockout and phenotypic characterization of mutants. (A) Schematic of *Bmmp* gene structure and nucleotides targeted for mutagenesis by CRISPR/Cas9. Genomic targets (not including PAM) are shown in underlined text, and PAM sequences are shown in red. Black rectangles and broken lines represent exons and introns, respectively, and are not to scale. (B) *Bmmp* knockout mosaics (*mp* KO) in the injected generation (G_0_) exhibited naked third abdominal segments in the pupal stage, suggestive of a changed wing phenotype. (C) *Bmmp* knockout mosaics (*mp* KO) in G_0_ exhibited significantly smaller wing areas by visual examination. Scale bar = 10 mm. (D) Mutant alleles detected by sequencing in nine randomly selected G_1_ silkworms. Red, base insertion; bold, base substitution; small deletions are represented with dashes; for large deletions, the sizes of the deleted regions are shown in parentheses. (E) Representative photograph of wings of WT and homozygous or compound heterozygous *Bmmp* knockout silkworms. Note that scale hairs were removed from the wings to exhibit the wing shape characteristics more clearly. Scale bar =2.5 mm (F) Measurement of total wing areas in WT and homozygous or compound heterozygous *Bmmp* knockout silkworms. N=23 for WT and N=45 for *Bmmp* knockouts. WT, wildtype control, *Dazao*; *mp* KO, homozygous or compound heterozygous *Bmmp* knockout silkworms. ***, P < 0.001.

To further confirm the function of *Bmmp* in a uniform genetic background, homozygous or compound mutant silkworms were obtained by crossing mosaic knockouts. We randomly surveyed 3 egg batches of generation 1 (G_1_). All individuals surveyed were homozygous or compound mutants (**Fig 5D**), demonstrating that germline transmission of the mutations was highly efficient. Compared with the WT control, homozygous or compound mutant silkworms all exhibited significantly smaller wings (**Fig 5E and 5F**), consistent with the phenotype observed in G_0_ mosaics.

It is noteworthy that we obtained homozygous knockout silkworms with an inframe 108 bp deletion in the coding region of exon 1 by crossing G_1_-mp-24♀ with G_1_-mp-15♂ (**Fig 5D)**. Presumably this mutation disrupts the functional BTB domain without affecting the downstream BACK and TLDc domains (**Fig 5D)**. However, these knockout silkworms were identical in wing phenotype to silkworms harboring frameshift mutations that presumably cause premature termination and functional loss of all three domains. These results suggest that the BTB domain plays an indispensable role in *Bmmp* gene function.

Taken together, we conclude that *Bmmp* plays an important role in wing development and the regulation of wing shape. Loss of function of *Bmmp* results in significantly smaller wings. In addition, we found that the BTB domain is indispensable for *Bmmp* function. We next sought to identify the mechanism(s) by which *Bmmp* regulates wing morphology.

### *Bmmp* regulates genes responsible for wing A-P and P-D axis development

The decreased wing size of *Bmmp* biallelic knockout silkworms reflects decreases in wing width and length along the anterior-posterior (A-P) and proximal-distal (P-D) axes, respectively. To detect potential interactions between *Bmmp* and other genes involved in wing formation, we used qRT-PCR to investigate the expression of key genes responsible for wing A-P and P-D axis development in wing discs from wandering stage *Bmmp* knockout silkworms. mRNA levels were significantly decreased in *Bmmp* knockout homozygous and compound heterozygous silkworms for *engrailed*, *hedgehog*, *dpp*, and *gbb,* which are responsible for wing A-P axis development (**Fig 6**). Likewise, mRNA levels in knockouts were reduced for *apterous A*, *apterous B*, *vestigial*, *Wingless* (*wnt1*), and *distal-less*, which participate in wing P-D axis development (**Fig 6**). These results suggest that *Bmmp* directs wing morphology by regulating genes responsible for wing A-P and P-D axis development.

**Figure 6.**
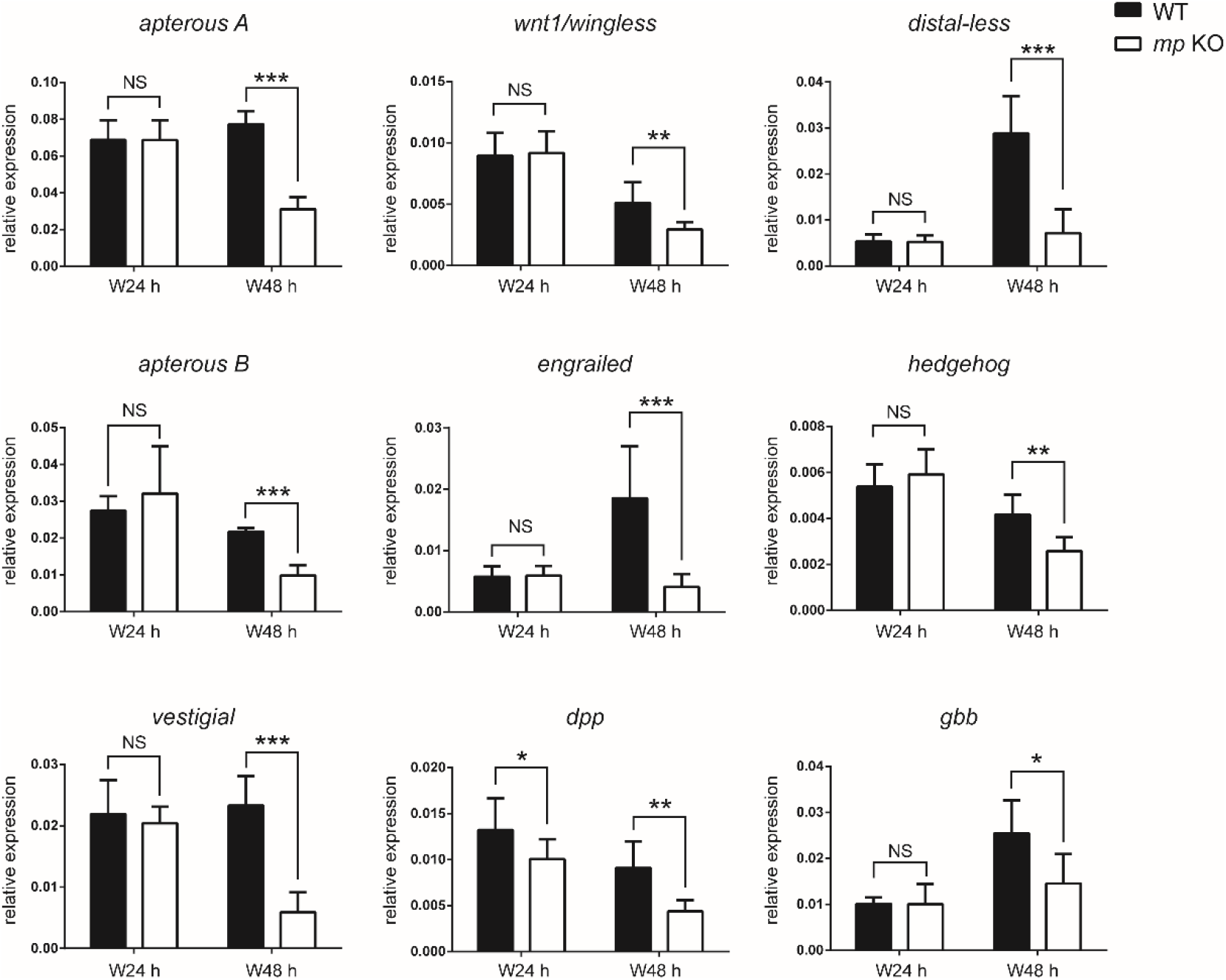
Expression of key genes responsible for wing A-P and P-D axis development in wandering stage silkworms. mRNA levels for *engrailed*, *hedgehog*, *wnt1*/*wingless*, *dpp*, *gbb*, *apterous A*, *apterous B*, *vestigial*, and *distal-less* were measured by qRT-PCR in wing discs from WT and *mp* silkworms at 24 (W24 h) and 48 hours (W48 h) after initiation of the wandering stage. NS, not significant; *, P < 0.05; **, P < 0.01; ***, *P* < 0.001; N=5 or 6 for *Dazao* and N=10 for *Bmmp* knockouts at 24 h; N=5 or 6 for *Dazao* and N=5 or 8 for *Bmmp* knockouts at 48 h. Values are relative to expression of eukaryotic translation initiation factor 4A (defined as 1). Experiments were independently repeated three times. WT, wildtype control, *Dazao*. *mp* KO, *Bmmp* knockout homozygous or compound heterozygous silkworms.

### *Bmmp* gene function is conserved between silkworms and *Drosophila*

The orthologous *Bmmp* gene in *Drosophila* is *CG7102*. Downregulation of this gene protects *Drosophila* from hypoxic tissue injury [16]. However, the functional role for *CG7102* in *Drosophila* wing development is not known. To examine whether the function of these two genes is conserved, we first predicted the functional domains of the *CG7102* protein product using the SMART online tool. Like *Bmmp*, *CG7102* is predicted to encode BTB, BACK, and TLDc domains. We obtained a *Drosophila mp* mutant strain that contains an insertion-associated gene mutation in *CG7102* from the Bloomington Drosophila Stock Center. Compared to the *Drosophila yw* control, the wings of the *Drosophila mp* mutant are curly and significantly smaller in total wing area, although the size difference between the *Drosophila* strains is not as severe as for *Bmmp* knockout and WT silkworms (**Fig 7A and 7B**). We speculate the milder phenotype may be due to genetic differences as the *Dropsophila mp* mutant contains an intronic transposon insertion in the *CG7102* gene, whereas the silkworm *Bmmp* knockouts we generated disrupt exon 1. Our tests show that the *Drosophila mp* mutant suffers a complete loss of flight capacity (**Video S1**), while flight is normal in the WT control (*yw*).

**Figure 7.**
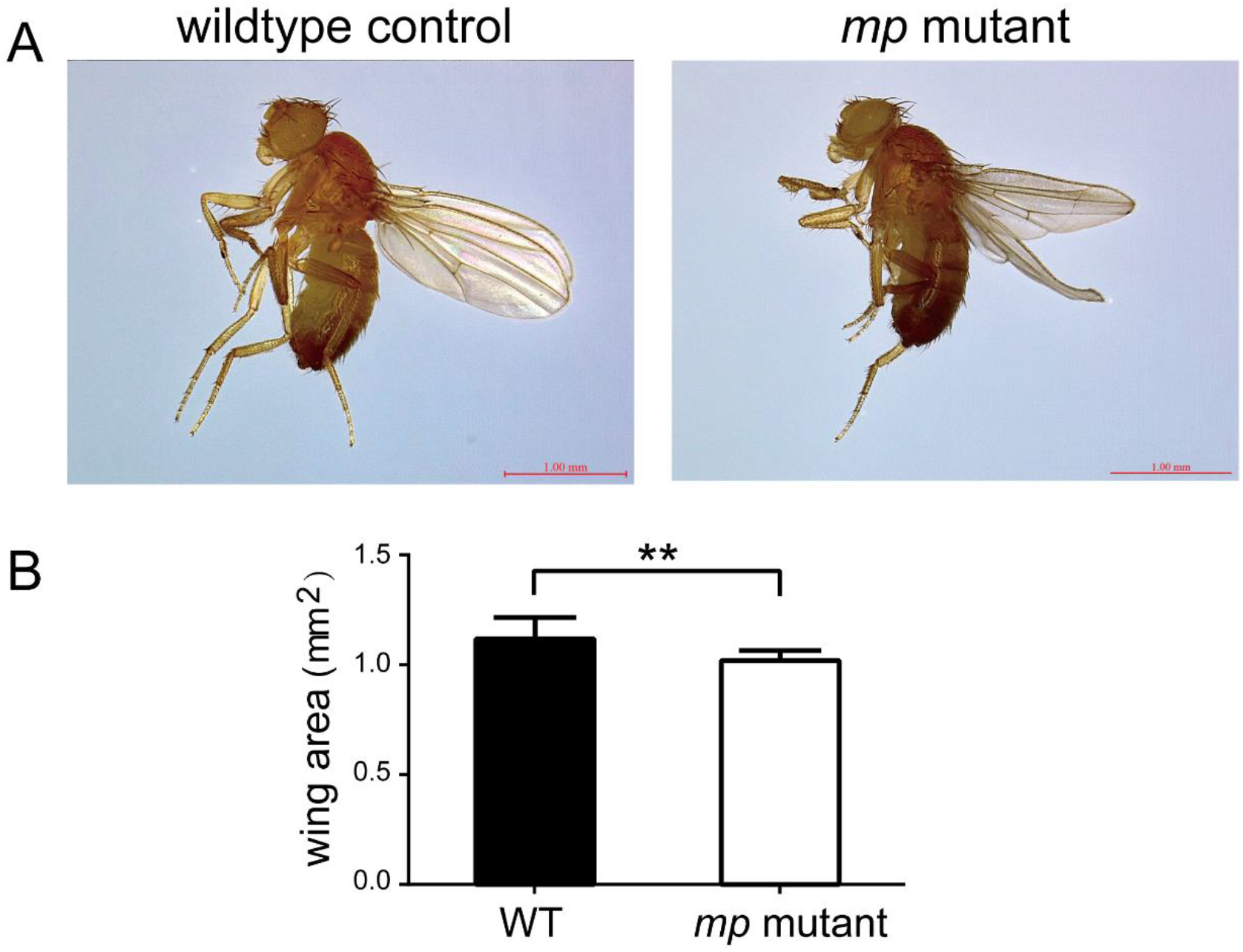
Wing phenotype of the *Drosophila mp* mutant. (A) The *Drosophila mp* mutant exhibited curly wings as compared with the WT control (*yw*) by visual examination. Scale bar, 1 mm. (B) Mean total wing area measurements. *Drosophila mp* mutants had significantly smaller wing areas compared with the WT control (*yw*). *, P < 0.01. N = 16 for *yw Drosophila* and N = 13 for *mp* mutants. *yw Drosophila* and *mp* mutants were both male. Experiments were repeated three times independently. Error bars represent SD.

## Discussion

The silkworm *mp* mutant exhibits shortened wing length and enlarged spacing of adjacent longitudinal veins without a decrease in total wing area. In this study, we identified the gene responsible for the *mp* phenotype and designated it *Bmmp*. Two possible causes for the *mp* phenotype are (1) a significant decrease in total *Bmmp* mRNA levels, and (2) the reduced diversity of *Bmmp* transcript isoforms in the wing discs. The changes in *Bmmp* expression in the silkworm *mp* mutant are likely to be caused by one or more mutations dispersed in non-coding regions of this gene. However, additional experiments would be required to dissect the effects of each of the non-coding mutations. In contrast, frameshift mutations induced by CRISPR/Cas9 mutagenesis into the constitutive exon 1 coding region of *Bmmp* resulted in significantly decreased wing length, width, and total wing area.

Alternative splicing is a ubiquitous regulatory mechanism of gene expression in eukaryotic organisms. For example, 90% to 95% of human genes are estimated to undergo alternative splicing [17, 18]. Variable mRNA transcript isoforms are translated into different protein isoforms with diverse functions and/or localizations [19]. In our study, we detected 28 distinct *Bmmp* transcript isoforms in wing discs of WT silkworms. Our findings suggest that *Bmmp* exerts its effect on wing development, at least in part, by exploiting diversified transcript isoforms, which give rise to different protein products with varying combinations of functional domains. Bmmp proteins can be categorized into three classes, namely, BTB-BACK-TLDc containing proteins, BTB-BACK containing proteins, and BTB-only proteins. Our results provide insight into the function of the BTB domain. A homozygous knockout silkworm harboring a 108-bp deletion that only disrupts the BTB domain exhibited the same wing phenotype as knockouts harboring frameshift mutations that presumably cause loss of function for all three domains. These results suggest the BTB domain is essential and indispensable for the function of the Bmmp protein. However, further investigation is needed to fully understand the functions of the BTB, BACK, and TLDc domains.

Pest migration, which depends on strong flight performance, is one of the most significant causes of damage to crops and forests [20]. Wing shape has a significant impact on the flight capacity of insects [1, 2]. Therefore, key genes regulating wing shape development are potential targets for pest control [21, 22]. In this study, we demonstrated that the *mp* orthologous gene *CG7102* is functionally conserved in *Drosophila*. Furthermore, we found flight capacity is completely lost in the *Drosophila mp* mutant, in which CG7102 contains a transposon insertion. Since mutations responsible for the *mp* phenotype compromise flight ability, it may be possible to exploit them in future pest control strategies. For example, if mutants can be released to cross into and reduce the fitness of target populations, the use of broad-spectrum pesticides could be reduced or avoided.

Finally, we demonstrated that null mutations of *Bmmp* decreased mRNA levels for genes involved in wing A-P axis development, including *engrailed*, *hedgehog*, *Dpp*, and *Gbb*, as well as genes involved in P-D axis development, including *apterous A*, *apterous B*, *vestigial*, *Wnt1*, and *Distal-less*. These results indicate that the *Bmmp* gene influences wing shape and size by regulating A-P and P-D axis development.

In summary, our results deepen our understanding of wing development in insects and provide a foundation for the development of insect pest control strategies.

## Materials and Methods

### Silkworm and *Drosophila* strains

Silkworm *Dazao* (wildtype) and *mp* mutant strains were obtained from the Silkworm Gene Bank at Southwest University (Chongqing, China). Silkworms were reared on fresh mulberry leaves at 25°C.

A *Drosophila melanogaster mp* mutant was purchased from the Bloomington Drosophila Stock Center (Stock Number: 80643). *Drosophila melanogaster* strain *yw,* with the same genetic background as the *Drosophila mp* mutant, was obtained from SIBCB Drosophila Library (Shanghai, China) and used as wildtype control in all *Drosophila* experiments. *Drosophila* strains were maintained at 25 ̊C with standard corn meal medium.

### Wing morphological measurements

Wings were dissected from silkworms (*Dazao*, *mp* mutant, *Bmmp* knockout mutant) and *Drosophila* (*yw*, *mp* mutant). Wings were imaged using a Leica DVM6 digital microscope. Wing shape parameters (wing area, wing length, adjacent vein spacings) were measured on 30 male and 30 female WT and *mp* silkworm moths, 23 female WT and 45 female *Bmmp* knockouts, and 16 male *yw* and 13 male *mp Drosophila* using ImageJ [23]. Experiments were independently repeated three times.

### Positional cloning and molecular mapping

*Dazao* and *mp* silkworms served as parental strains to produce F_1_ progeny. Due to the lack of recombination in female silkworms, 20 progeny from a single-pair backcross between an F_1_ female and a *mp* male (BC_1_F) were used for the linkage analysis and 456 progeny from *mp* female × F_1_ male backcrosses (BC_1_M) were used for the recombination analysis. Developing embryos were incubated at 25°C in a humidified atmosphere.

We performed preliminary mapping using published SSR markers [24, 25]. SSR markers on chromosome 11 that were polymorphic between the parental strains were used for linkage and recombination analyses. Linkage analysis was conducted with JOINMAP 4.0 using Kosambi’s mapping function [26]. SSR markers were used as anchor points to develop novel markers based on the silkworm genome sequence (International Silkworm Genome Consortium, 2008) for fine mapping with 320 BC_1_M.

### Identification of *mp*-specific SNPs and indels

Silkworm *mp* mutants were analyzed by whole genome sequencing according to a previously published protocol [27]. To identify *mp-*specific mutations, data were compared to sequences from 127 domestic and wild silkworm strains with normal wing phenotypes from SilkBase [15] and a previous report [27]. Alignments to reference sequences (released in November 2016 by SilkBase [15]) were performed to identify SNPs and indels in the *mp* mutant and the 127 silkworm strains, respectively. SNPs and indels were extracted from genomic sequences surrounding *KWMTBOMO06923* and *KWMTBOMO06924* in the *mp* strain and the 127 silkworm strains, and then screened for variations specific to *mp* mutants using an online Venn diagram tool (http://bioinformatics.psb.ugent.be/webtools/Venn/).

### Examination of *Bmmp* transcript isoforms

To generate *Bmmp* cDNA libraries, total RNA was extracted from wing discs of *mp* and WT silkworms. For each strain, wing disc samples were obtained from silkworms from fifth-instar larvae on days 3, 5, and 7 (5L3D, 5L5D, and 5L7D) and from wandering stage silkworms at 0, 24, and 48 hours (W 0 h, W 24 h, and W 48 h). RNA extractions were performed using the MicroElute Total RNA Kit (OMEGA), and reverse transcription was performed using the PrimeScript RT Reagent Kit (Takara). Equal masses (concentration times volume) of the resulting cDNAs from different developmental stages were mixed and used as the templates for PCR amplification. The PCR primers were F: 5’-AAACTAAACTTATTTGAGGTTATG-3’, and R: 5’-AATAATCATCGGACTAAATCACCTT-3’. PCR products were subcloned into pEASY-blunt-zero vectors (TransGen) and sequenced.

### Quantitative RT-PCR (qRT-PCR)

Total RNA was extracted from wing discs of individual silkworms at the wandering stage using the MicroElute Total RNA Kit (OMEGA) and reverse transcription was performed using the PrimeScript RT reagent Kit (Takara). qRT-PCR experiments were performed using Hieff SYBR Green Master Mix (YEASEN), according to the manufacturer’s recommended procedure. Silkworms were sampled as follows: N=5 or 6 for *Dazao* and N=10 for *Bmmp* knockouts at 24 h of the wandering stage; N=5 or 6 for *Dazao* and N=5 or 8 for *Bmmp* knockouts at 48 h of the wandering stage. Three independent replicates were performed for all qRT-PCR experiments. Primer sets are listed in **Table S3**. Eukaryotic translation initiation factor 4A (silkworm microarray probe ID sw22934) was used as the internal control.

### *Bmmp* knockout generation

sgRNAs for CRISPR/Cas9 mutagenesis were designed using the CHOP-CHOP online utility (http://chopchop.cbu.uib.no/). sgRNA target sites are shown in Figure 4A. The DNA template for the T7 promoter used to drive *in vitro* transcription was constructed by PCR as described [28]. Briefly, an oligonucleotide containing the T7 promoter and the sgRNA target sequence (N_20_) was designed as a forward primer with the sequence 5’-<u>TAATACGACTCACTATAGG</u>(N_20_)GTTTTAGAGCTAGAAATAGC. The T7 promoter sequence is underlined. The reverse primer was 5’-AAAAGCACCGACTCGGTGCCACTTTTTCAAGTTGATAACGGACTAGCCTTA TTTTAACTTGCTATTTCTAGCTCTAAAAC-3’. sgRNA synthesis was performed using a T7 RiboMax Large Scale RNA Production System (Promega) following the manufacturer’s instructions.

The bivoltine silkworm strain *Dazao* was used to generate *Bmmp* knockout silkworms. To generate non-diapaused eggs, silkworm eggs were incubated at 15°C until hatching, and the larvae were reared on fresh mulberry leaves at 25°C until wandering stage. Adult moths then oviposited non-diapaused eggs, which were used for microinjection. A mixture of sgRNA and Cas9 protein (Thermo Fisher) was incubated at room temperature for 15 min and microinjected into preblastoderm embryos within 5 h of oviposition. Injected embryos were incubated at 25°C and 80% humidity for approximately 10 days until hatching. Larvae were maintained at 25°C and fed fresh mulberry leaves.

### Identification of *Bmmp* knockout silkworm genotypes

Genomic DNA was extracted from the wings of *Bmmp* knockout silkworms at the adult stage using the TIANamp Genomic DNA Kit (TIANGEN). The DNA was used as a PCR template to amplify regions spanning the genomic targets. Two primer sets were used as follows. F1: 5’-TCGGAGCCGTCTTTAAGTGT-3’ and R1: 5’-CAGAAGATGGTTAAGATGACGTT-3’, and F2: 5’-GGTTGCGTTGGTGGTGTAAT-3’ and R2: 5’-TTATCCTGCCCAGCTGAGAG-3’. PCR products were subcloned into pEASY-blunt-zero vectors (TransGen), and sequenced.

### Statistical analysis

All values are presented as means ± SEM or means ± SD, as indicated in figure legends. Student’s t test was used to determine p values.

## Abbreviations

A-P: anterior-posterior
cDNA: complementary DNA
CRISPR/Cas9: clustered regularly interspaced short palindromic repeats/CRISPR-associated protein-9 nuclease
D-V: dorsal-ventral
*mp*: *micropterous*
P-D: proximal-distal
qRT-PCR: quantitative real-time PCR
SNP: single nucleotide polymorphism
SSR: simple sequence repeat
WT: wildtype.

## Acknowledgments

We wish to thank our group members for their continuous support.

## Data Availability Statement

All relevant data are within the manuscript and its Supporting Information files.

